# Mutual regulation of transcriptomes between pneumocytes and fibroblasts mediates alveolar regeneration

**DOI:** 10.1101/2023.02.27.530149

**Authors:** Yiwen Yao, Sarah Miethe, Kathrin Kattler, Jörn Walter, Nicole Schneider-Daum, Christian Herr, Holger Garn, Felix Ritzmann, Robert Bals, Christoph Beisswenger

**Affiliations:** Department of Internal Medicine V – Pulmonology, Allergology and Critical Care Medicine, Saarland University, 66421 Homburg, Germany; Department of Clinical Medicine, Shanghai Tongji Hospital, School of Medicine, Tongji University, 200065 Shanghai, China; Translational Inflammation Research Division & Core Facility for Single Cell Multiomics, Member of the German Center for Lung Research (DZL) and the Universities of Giessen and Marburg Lung Center, Medical Faculty, Philipps University of Marburg, D-35043 Marburg, Germany; Department of Genetics/Epigenetics, Saarland University, 66123 Saarbrücken, Germany; Helmholtz Institute for Pharmaceutical Research Saarland (HIPS), Helmholtz Centre for Infection Research (HZI), Saarland University Campus, 66123 Saarbrücken, Germany; Department of Drug Delivery (DDEL), Helmholtz-Institute for Pharmaceutical Research Saarland (HIPS), Helmholtz Centre for Infection Research (HZI), Saarbrücken, Germany

**Keywords:** alveolar regeneration, pneumocytes, fibroblasts, differentiation, IL-6, single-cell

## Abstract

Alveolar type 2 (AT2) and club cells are part of the stem cell niche of the lung and their differentiation is required for pulmonary homeostasis and tissue regeneration. A disturbed crosstalk between fibroblasts and epithelial cells contributes to the loss of lung structure in chronic lung diseases. Therefore, it is important to understand how fibroblasts and lung epithelial cells interact during regeneration. Here we analyzed the interaction of fibroblasts and the alveolar epithelium modelled in air-liquid interface cultures. Single-cell transcriptomics showed that co-cultivation with fibroblasts leads to increased expression of type 2 markers in pneumocytes, activation of regulons associated with maintenance of alveolar type 2 cells, and trans-differentiation of club cells towards pneumocytes. This was accompanied by an intensified transepithelial barrier. Vice versa, activation of NFκB pathways and the CEBPB regulon as well as the expression of IL-6 and other differentiation factors (e.g. FGFs) were increased in fibroblasts co-cultured with epithelial cells. Recombinant IL-6 enhanced epithelial barrier formation. Therefore, in our co-culture model, regulatory loops were identified by which lung epithelial cells mediate regeneration and differentiation of the alveolar epithelium in a cooperative manner with the mesenchymal compartment.

## Introduction

The epithelial surfaces of the lung extend from the upper airways to alveoli. The airway epithelium consists of a pseudostratified ciliated epithelium composed primarily of basal cells, mucus-secreting goblet cells, club cells, and ciliated cells. In the alveoli, flat alveolar type 1 (AT1) cells constitute most of the alveolar surface for gas exchange, while cubic AT2 cells ensure normal lung function by secreting various factors such as surface tension-lowering surfactant [1].

The maintenance and coordinated differentiation of different cell types with stem cell capacity is essential for lung homeostasis and regeneration. Single cell analyzes showed that basal cells represent the progenitor cells for club, goblet and ciliated cells, with the latter two differentiating via club cells [2, 3]. In the alveolar region, self-renewing type II pneumocytes are progenitors of AT1 cells. AT2 cells therefore have an important function in homeostasis and regeneration of the lung parenchyma after damage caused, for example, by infection [4]. Mouse studies suggest that club cells located at the terminal bronchioles serve as progenitor cells not only for airway cells but also for AT2-like cells and are likely involved in regeneration of the lung after severe injury. Whether these club cells, which express AT2 markers, actually differentiate into true AT2 cells, or whether they have other, unknown functions in the regeneration and repair of the lung parenchyma after severe damage, is not completely understood [5–9]. Single cell analysis revealed that the human respiratory bronchioles also contain secretory airway epithelial cells that are able to differentiate into AT2 cells [10].

Fibroblasts express components of the extracellular matrix (e.g. collagen, fibronectin) and interact closely with epithelial cells in the regulation of proliferation and differentiation. Fibroblasts are dynamic cells that can differentiate into different, overlapping subtypes comprising peribronchial fibroblasts, adventitial fibroblasts, myofibroblasts, and alveolar fibroblasts [11, 12]. Various studies have addressed how mesenchymal cells regulate the proliferation and differentiation of lung epithelial cells and, conversely, how epithelial cells affect fibroblasts. This epithelial-fibroblast crosstalk is critical for lung development, maturation, homeostasis and repair after injury and numerous signaling pathways such as wingless-related integration site (WNT) and fibroblast growth factors (FGF) are involved [12, 13]. Lee et al. showed, for example, using mouse and organoid studies, that mesenchymal cells promote the differentiation of airway progenitor cells via a Wnt-Fgf10 cooperation. Mesenchymal cells also mediate alveolar differentiation of progenitor cells in a WNT-dependent manner [14]. Zepp al. showed that diverse mesenchymal lineages mediate the growth and self-renewal of AT2 progenitor cells via IL-6/Stat3 and Fgf signaling in mice [15]. The aim of the present study was to analyze the interaction of fibroblasts and lung alveolar cells by single-cell RNA sequencing (scRNA-seq). We show that isolated AT2 cells cultured in air-liquid interface cultures differentiate towards AT1 cells. The coculture with fibroblasts leads to an increased maintenance of type 2 cells and transdifferentiation of club cells to AT2-like cells. Conversely, epithelial cells induce increased expression of growth factors in fibroblasts.

## Results

### Air-Liquid interface (ALI) cultures resemble native alveolar cell composition

First, we characterized distal murine lung cells cultured in ALIs by scRNA sequencing. Mouse lung tissue was dissociated and EpCAM^+^ cells were isolated by MACS. The obtained cells were seeded in Transwells and the apical chamber medium was aspirated after 2 days. Single cell suspensions were obtained after 2 days post airlift and single cell transcriptome profiling was performed by BD Rhapsody™ Single-Cell Analysis System (Fig. 1A). Representative markers [11] were used to differentiate major cell types: epithelial (82.7%), macrophages (7.5%) and mesenchymal (9.8%) cells (Fig 1B, S1).

**Fig. 1.**
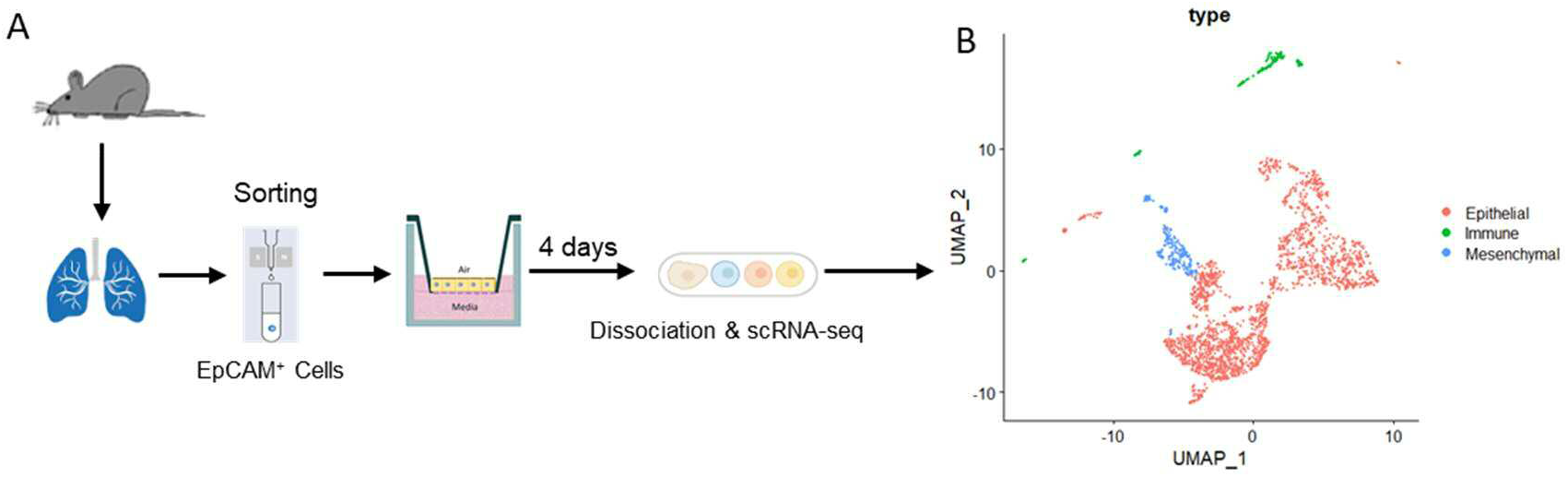
scRNA-seq analysis analysis of murine lung cells cultured in ALI. **(A)** Schematic representation of the experimental setup. Mouse EpCAM^+^ cells were isolated and cultured om Transwells for four days. **(B)** UMAP visualization of major cell types.

For further characterization, we sub-clustered epithelial cells. Fig. 2A shows that both, pneumocytes and airway epithelial cells were present in our ALI model and that cells from these compartments could be clearly distinguished in the Uniform Manifold Approximation and Projection (UMAP). The pneumocytes formed a continuum from AT2-like cells with high expression of Type 2 markers (e.g. *Sftpc, Lyz2, Lamp3*) via immature AT1 (*Hopx*^high^/*Sftpc*^medium^) cells to AT1-like cells with low expression of Type 2 markers and increased expression of *Hopx, Aqp5* and *Krt7* (Fig. 2B and C). We also identified a transitional cluster that expressed epithelial and mesenchymal markers at low level. *Scgb1a1-expressing* club cells were the dominant cells among the airway epithelial cells, which also included *Krt5-* and *Trp63*-expressing basal cells and *Mki67*-expressing proliferating cells (Fig. 2B and C). Immunofluorescence (IF) staining for AT2 (SFTPC) and club cells markers (CCSP) on membranes of ALI confirmed the principle cell types identified by scRNA-seq (Fig 2D and E). These data show that ALI cultures of murine lung cells resemble the cellular composition of the lower respiratory tract tissue.

**Fig. 2.**
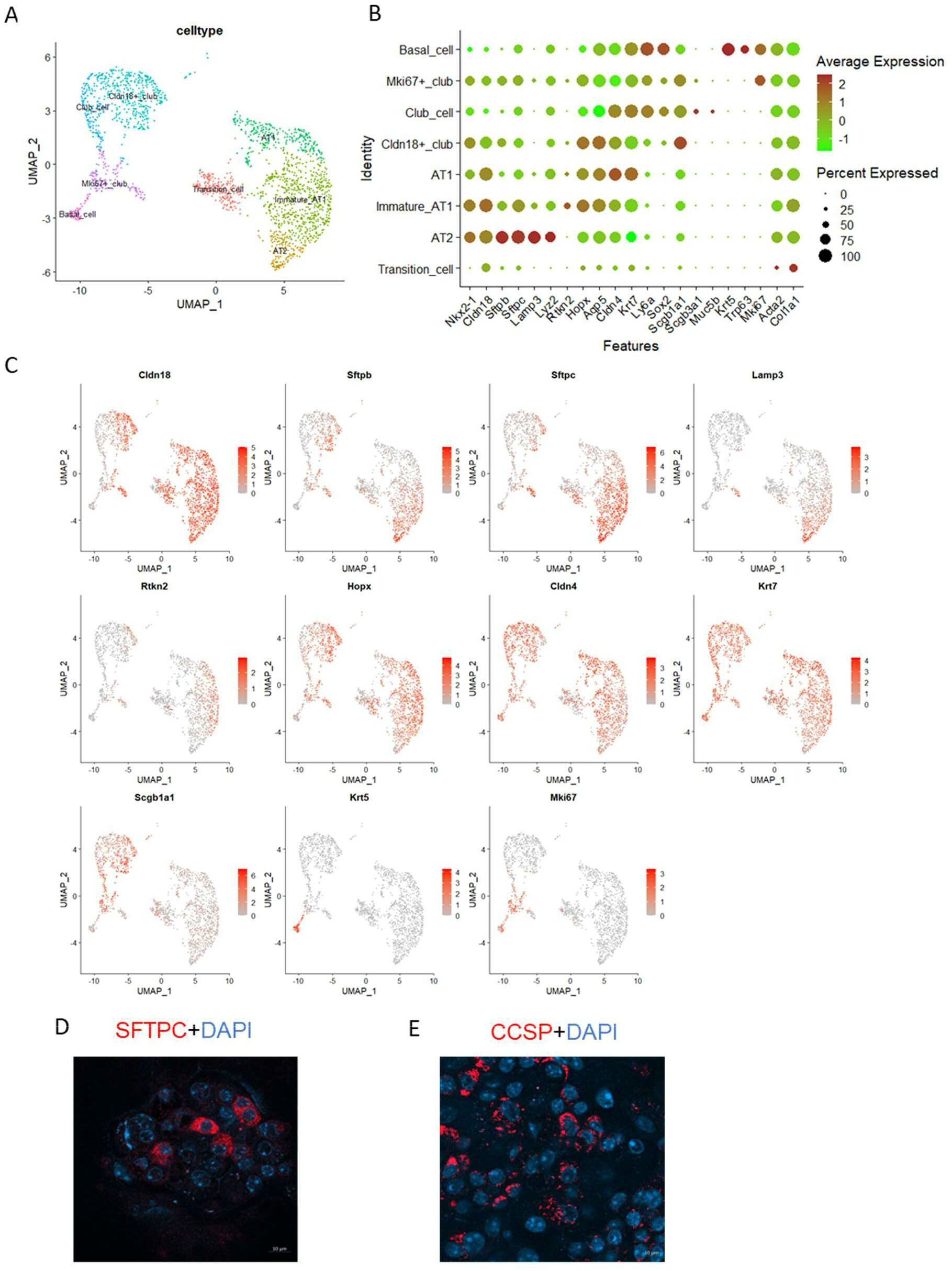
scRNA-seq analysis of ALI cultures reveals distinct lung epithelial cell populations. **(A)** UMAP visualization of major epithelial cell types. **(B)** Intensity dot plot showing expression of epithelial cell type markers. **(C)** UMAP visualization of cell type-specific markers expression. ALI-cultures were stained for **(D)** SFTPC and **(E)** CCSP by immunofluorescence.

### Fibroblasts promote AT-2 cell differentiation

In order to understand the role of fibroblasts in epithelial regeneration, we isolated EpCAM^-^ cells from mouse lungs and co-cultured the obtained fibroblasts with epithelial cells for four days (Fig. 3A). scRNA-seq analysis confirmed the expression of fibroblast markers (e.g. *Colla1, Acta2, Pdgfra*) in these mesenchymal cells at the end of the experiment (S2). We measured the transepithelial electrical resistance (TEER) of the ALI cultures to monitor whether the presence of fibroblasts affects epithelial barrier formation. Co-culture of epithelial cells with fibroblasts led to a significantly increased TEER after 48 hours of the ALI cultures (Fig. 3B).

**Fig. 3.**
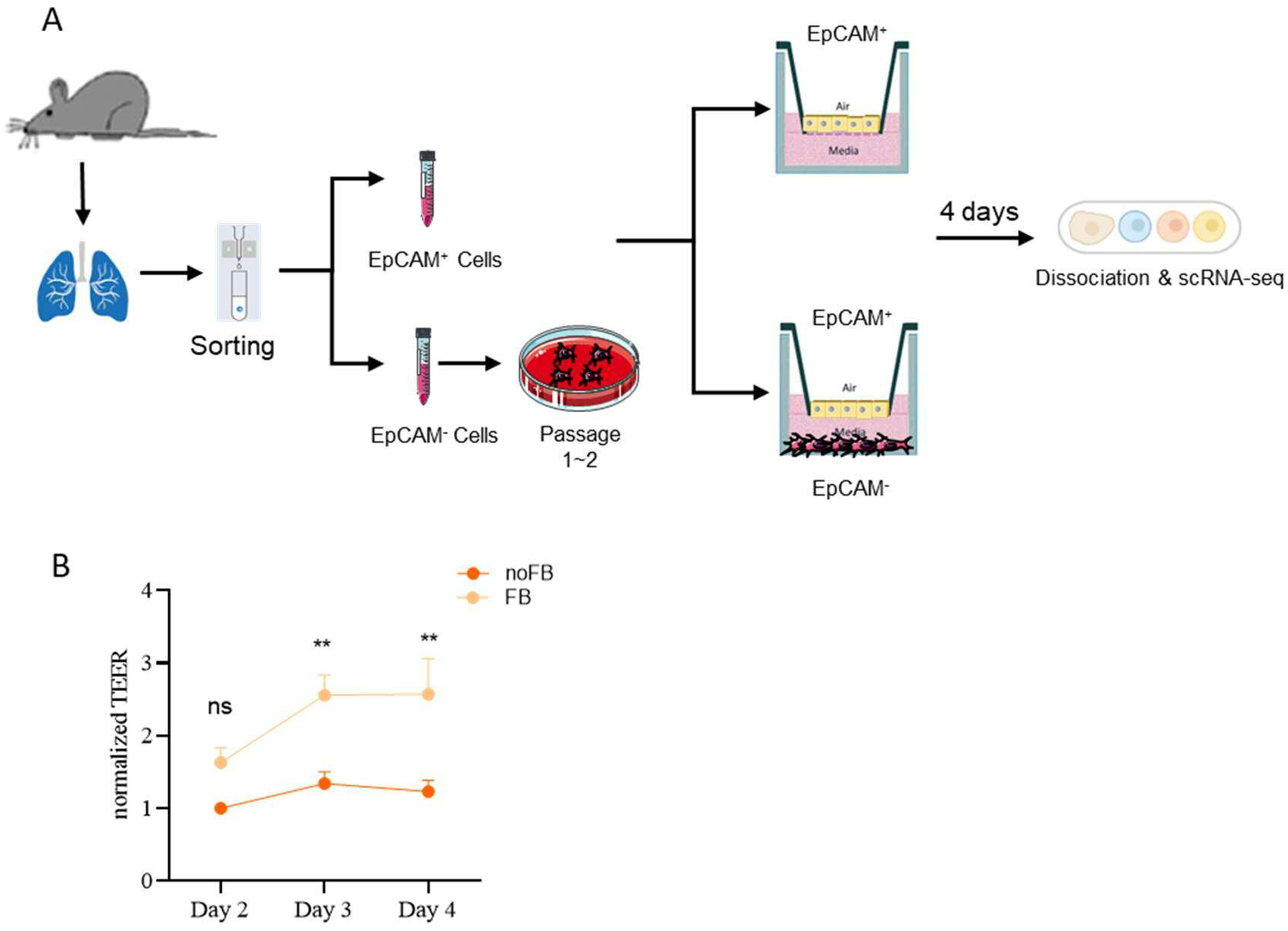
Fibroblasts promote the formation of epithelial barrier. **(A)** Schematic representation of the experimental setup. Mouse EpCAM^+^ cells were isolated and cultured in ALIs for four days with and without fibroblasts. **(B)** TEER was measured every 24 hours (n=5 independent experiments). Data were compared by two-way ANOVA. **p < 0.01.

scRNA-seq analysis of the epithelial cells showed that co-culture with fibroblasts resulted in an increased expression of AT2-markers and club cell proliferation. *Sftpb* (2.3-log2 fold change), *Cbr2* (1.8-log2 fold change) and *Cxcl15* (1.9-log2 fold change) as well as the club cell makers *Scgb1a1* (2.0-log2 fold change) and *Cyp2f2* (1.8-log2 fold change) were the top five up-regulated genes in the co-culture group (Fig. 4A). Figures 4B and C show that co-culture with fibroblasts specifically enhanced expression of AT2 markers in AT2 cells as well as in club cells, whereas the expression of the proliferation marker *Ccnd1* and the club cell markers *Scgb1a1* and *Cyp2f2* was only increased in club cells. qPCR analysis confirmed that co-culture with fibroblasts leads to an increased expression of pneumocyte and club cell markers in the ALI model (Fig. 4D). These data show that presence of fibroblasts results in the modification of epithelial differentiation towards AT2 cells.

**Fig. 4.**
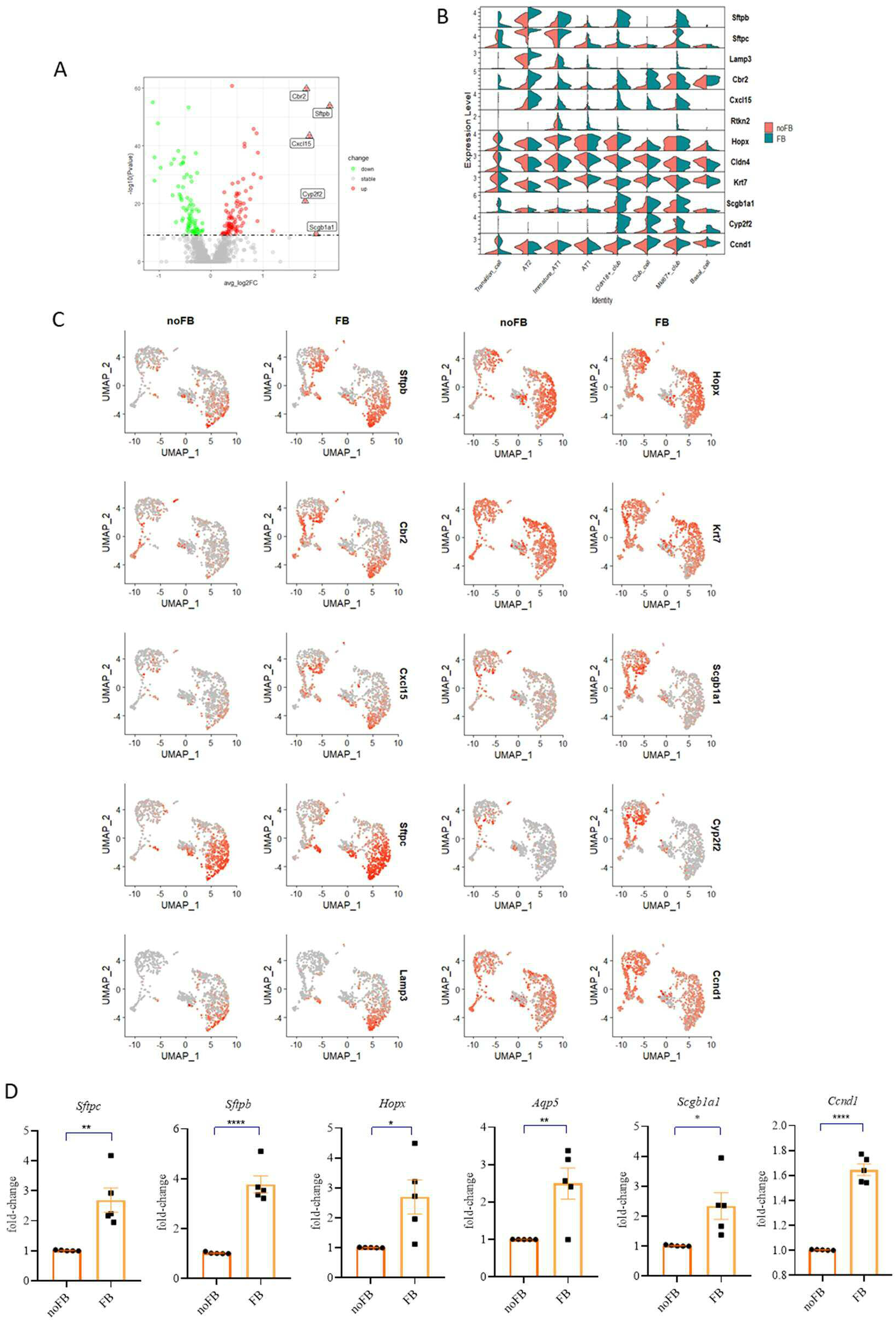
Fibroblasts enhance the expression of AT-2 markers. **(A)** DEG analysis of epithelial cells co-cultured with/without mesenchymal cells. **(B)** Violin plots and **(C)** feature plots showing the expression levels of differentially expressed cell markers. **(D)** The expression of selected cell markers was confirmed by semi-quantitative RT-PCR after 4 days culture. Data are shown as mean ± SEM (n=5 independent experiments). Data were compared by unpaired t test. *p < 0.05, **p < 0.01, ***p < 0.001, ****p < 0.0001.

Trajectory analysis was used to trace the differentiation pathways of AT2 and club cells. Mature AT2 cells differentiated to AT1 cells via immature *Hopx*^high^/*Sftpc*^medium^ AT1 cells. *Nkx2-1*^low^/*Cldn18*^low^ club cells differentiated to AT2 marker expressing *Cldn18*^high^/*Nkx2*^high^ club cells (Fig 5A).

**Fig. 5.**
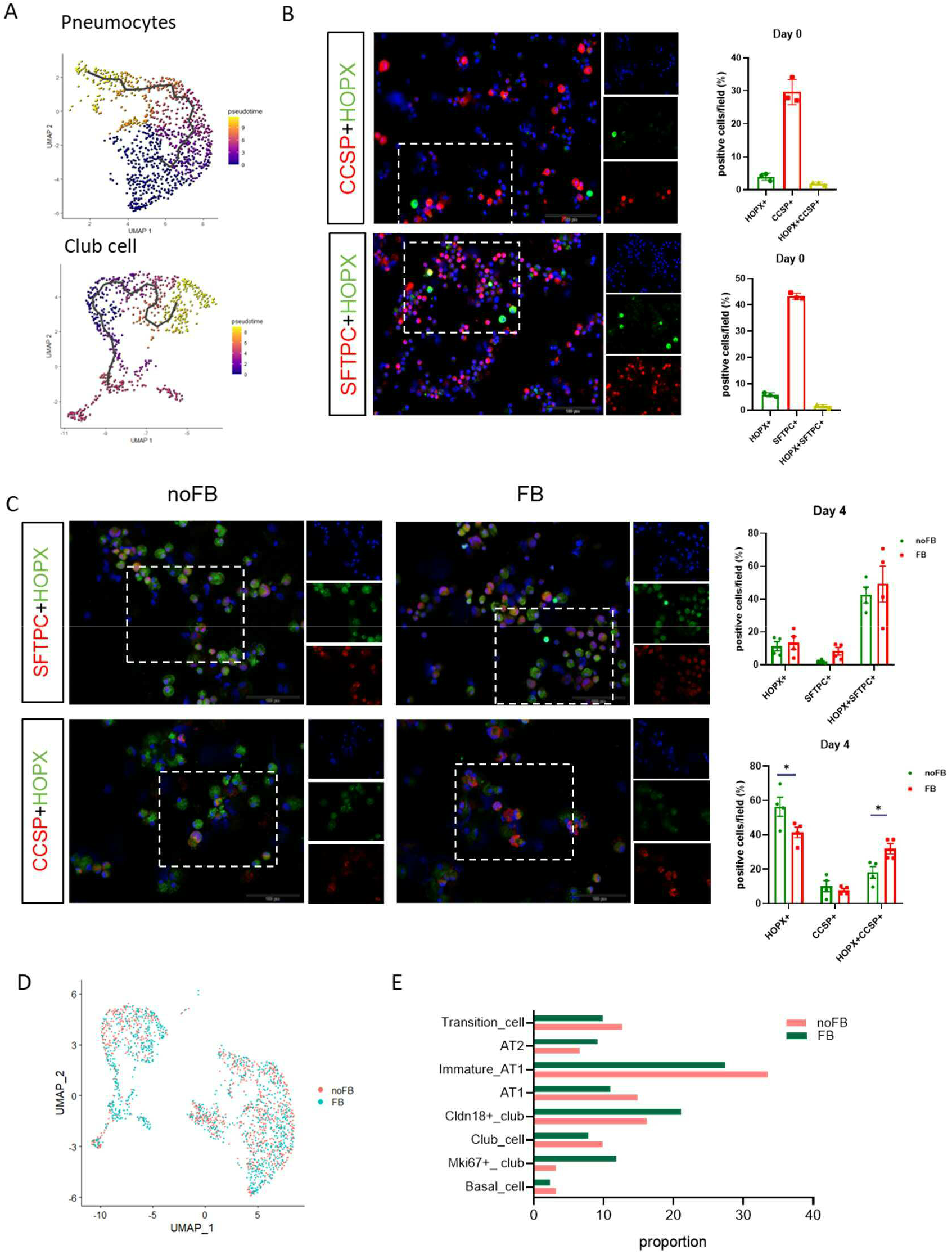
Trajectory analysis of epithelial cells. **(A)** Pseudotime trajectory of selected epithelial cell types from ALI cultures. Cytospins were prepared **(B)** immediately after isolation or **(C)** after four days of culture. SFTPC^+^, HOPX^+^, and CCSP^+^ were stained by immunofluorescence. The fraction of cells positive for the different markers was quantified. Data of a representative experiment. Data were compared by two-way ANOVA. *p < 0.05. **(D)** UMAP plot indicting epithelial cells with cocultured with fibroblasts (FB, blue) and without fibroblasts (noFB, red) **(E)** Relative proportion of different epithelial cell types in the presence or absence of fibroblasts.

To examine how the ratio of pneumocytes and club cells differs immediately after isolation and after four days in ALI culture, we analyzed cytospins of respective cultures for SFTPC, CCSP and HOPX by staining. IF showed that mainly AT2 cells (45% SFTPC^+^ cells) and club cells (30% CCSP^+^ cells) were seeded on the transwells immediately after isolation and that the ratio of AT2 cells to club cells hardly changed after 4 days of culture (Fig. 5 B and C). However, there was a significant increase in SFTPC^+^ cells and CCSP^+^ cells co-expressing HOPX. Increased expression of HOPX has been demonstrated during lung injury and repair [16, 17]. In co-culture with fibroblasts, the proportion of club cells expressing HOPX further increased. The scRNA-seq data also showed a similar ratio of the different cell types with increased numbers of *Cldn18^+^* club cells in the co-culture group (Fig 5. D and E). These results demonstrate a high ability of AT2 cells to differentiate towards AT1 cells.

### Fibroblasts activate regulons that mediate AT2 differentiation

To gain insight into transcriptional regulatory networks, we performed a regulon analysis using SCENIC [18] (Fig. 6A). The Irx1 regulon, which regulates AT2 cell differentiation [19], was found to be highly activated in pneumocytes independently of fibroblasts (Fig. 6 B and C). Etv5, which is essential for the maintenance of AT2 cells [20], showed increased activity in AT2 cells in cultures co-cultivated with fibroblasts (Fig. 6 D and E). We also found numerous regulons with unknown function, whose activity was increased in co-cultured club cells and AT2 cells (Fig. 6 F and G, S3). Six1, which has been repotted to play a role in the coordination of lung epithelial, mesenchymal, and vascular development [21], showed increased activity in club cells co-cultured with fibroblasts (Fig. 6 H and I).

**Fig. 6.**
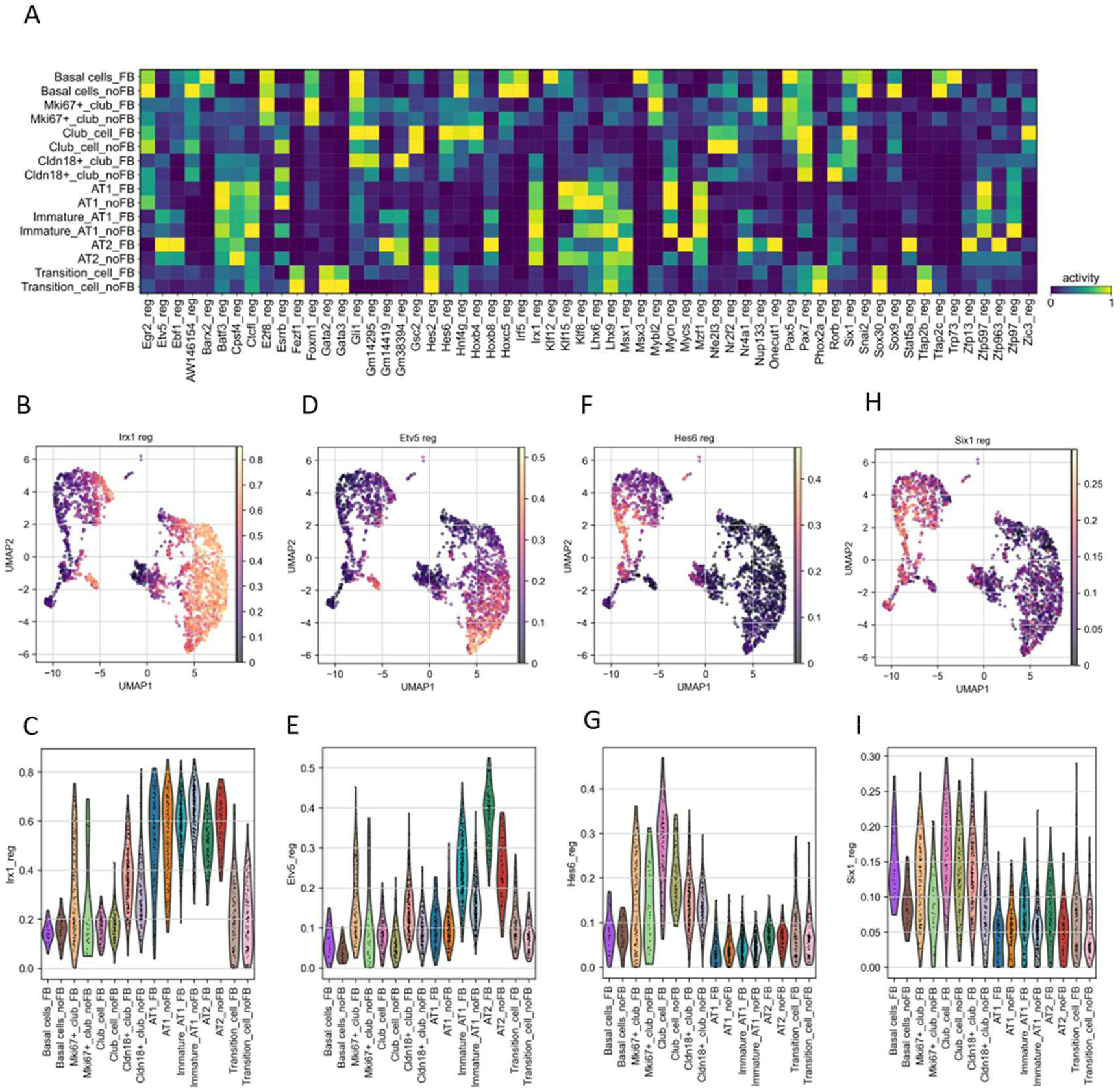
Regulon analysis in epithelial cells. **(A)** Heatmap of regulon activity of the top regulons discriminating FB and noFB clusters. UMAP representation and corresponding violin plots of regulon activity of (**B/C**) Irx1, (**D/E**) Etv5, (**F/G**) Hes6 and (**H/I**) Six1.

### Co-culture with epithelial cells leads to increased expression of growth factors in fibroblasts

We further analyzed the influence of epithelial cells on fibroblasts transcriptome profiles using scRNA-seq (Fig. 7A). Five cell clusters were identified within the population characterized by the expression of general fibroblast markers (e.g. *Col1a1*) (Fig. 7 B to D) of which four clusters showed robust expression of *Pdgfra. Pdgfra^+^* fibroblasts have been ascribed to play a role in alveolar proliferation and differentiation [22]. Two *Pdgfra* clusters were positive for *Mki67* and probably represent the proliferating fibroblast pool.

**Fig. 7.**
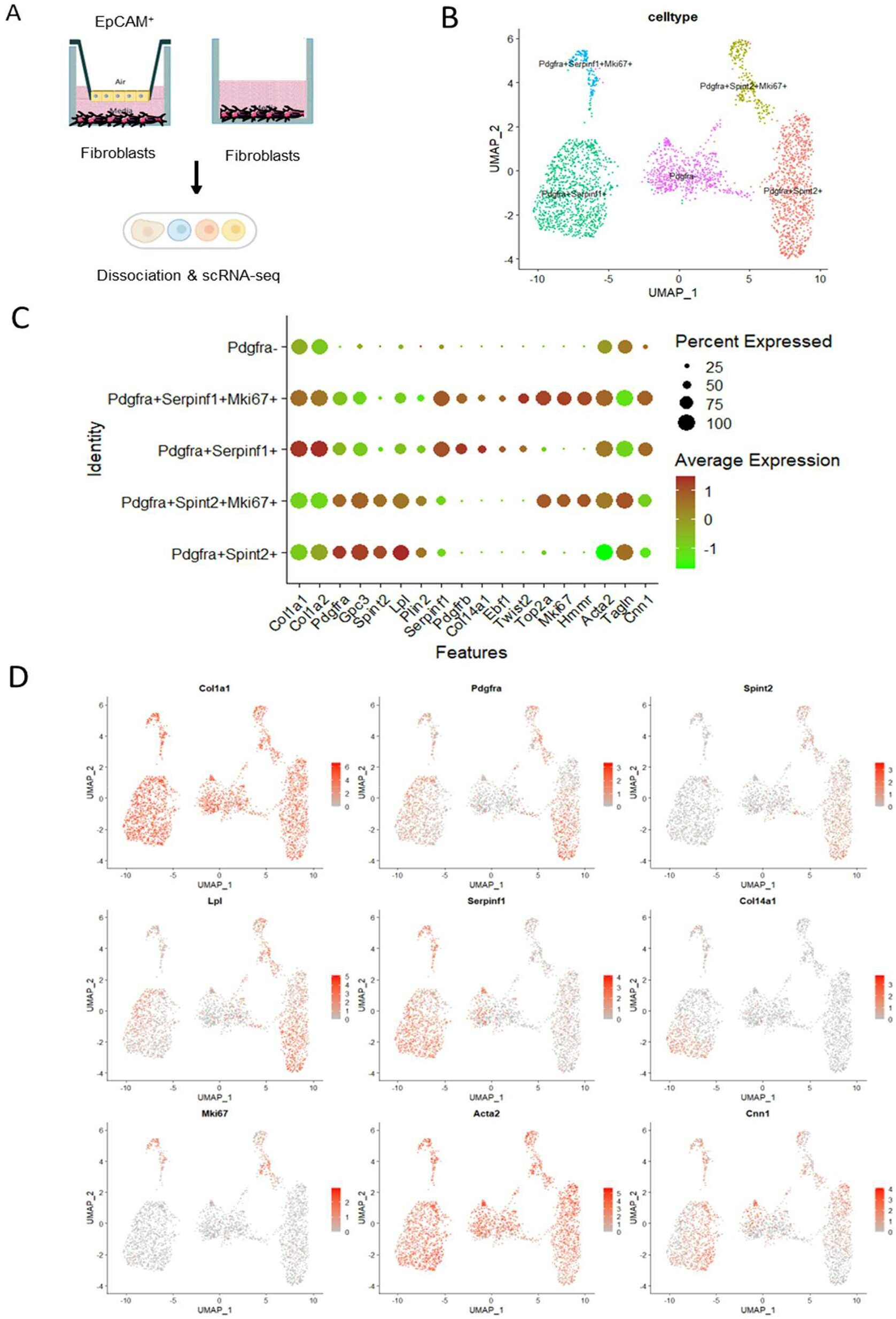
scRNA-seq analysis of fibroblasts. **(A)** Schematic for scRNA-seq of fibroblasts experiment. **(B)** UMAP visualization of fibroblasts. **(C)** Dot plot showing expression of fibroblasts type markers. **(D)** UMAP visualization of cell type-specific markers expression.

We compared fibroblasts co-cultured with or without epithelial cells (Fig. 8A). The proportion of the *Pdgfra^+^Serinf1^+^* cluster was increased by about 10 % after co-culture with epithelial cells (Fig. 8B). Growth factors associated with alveolar regeneration, such as FGF7, FGF10 and EREG [23], were highly expressed in *Pdgfra^+^Serinf1^+^* clusters. Co-culture with epithelial cell resulted in increased expression of these factors. (Fig.8 C and D).

**Fig. 8.**
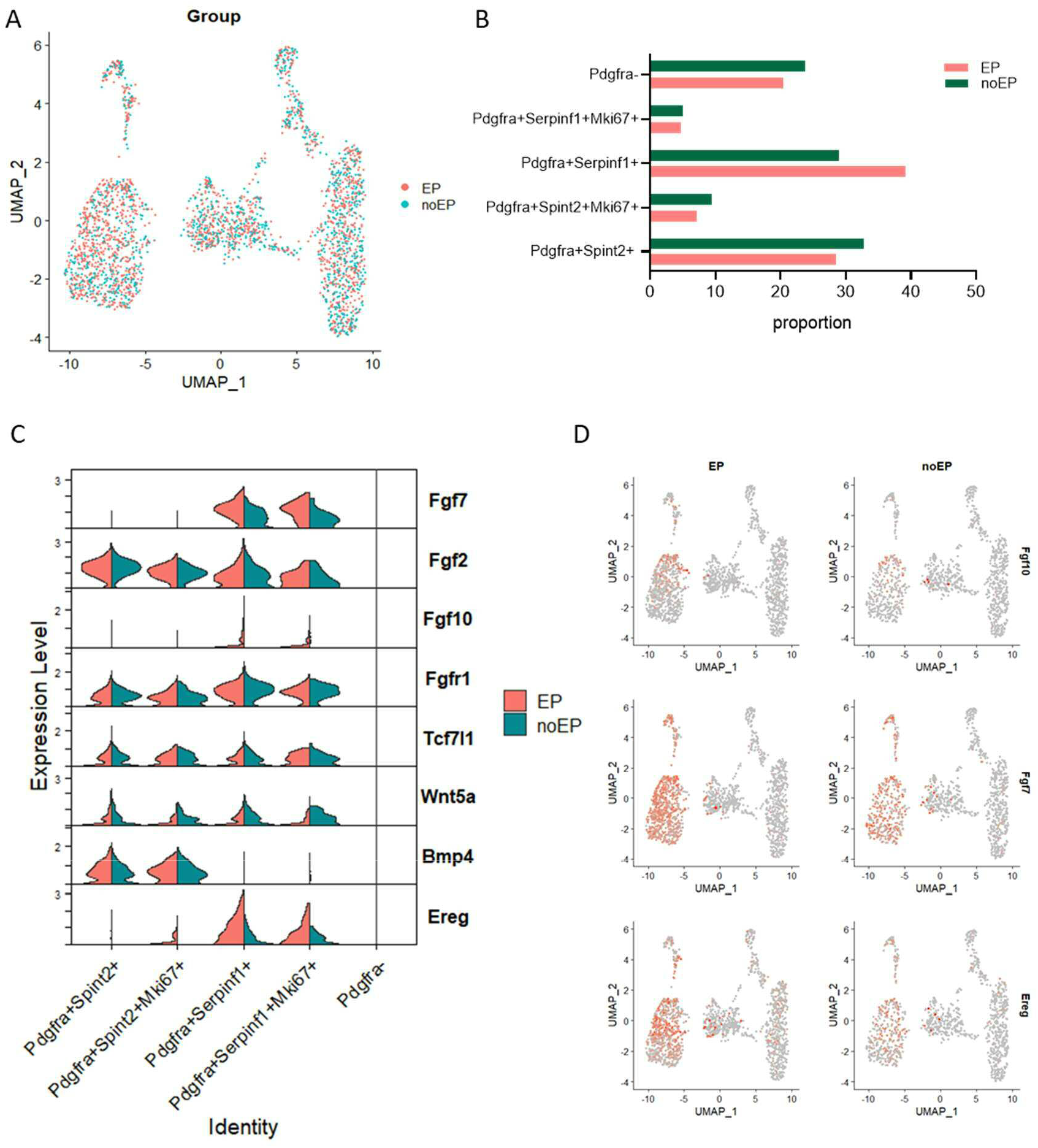
Epithelial cells induce the expression of growth factors in the *Pdgfra^+^Serinf1^+^* cluster. **(A)** UMAP plot indicating fibroblasts cultured in absence (noEP, blue) or presence of epithelial cells (EP, red). **(B)** Relative proportion of each fibroblasts subtype in noEP versus EP group. **(C)** Violin plots and **(D)** feature plots showing the expression levels of differential expressed fibroblast ligands.

### Co-culture with epithelium results in the activation of TNFA/NFKB and IL-6/STAT3 pathways in fibroblasts

Pathway analysis using gene set variation analysis (GSVA) and AUCell showed that TNFA/NFKB and IL-6/STAT3 pathways were dramatically activated when fibroblasts were co-cultured with epithelial cells, especially in *Pdgfra^+^Serinf1^+^* fibroblasts (Fig. 9 A and B). Co-culture with epithelial cells also increased the activity of Cebpb regulon in *Pdgfra^+^Serinf1^+^* fibroblasts (Fig. 9 C and D), which is important in the regulation of genes involved in immune and inflammatory responses and enhances IL-6 transcription [24].

**Fig. 9.**
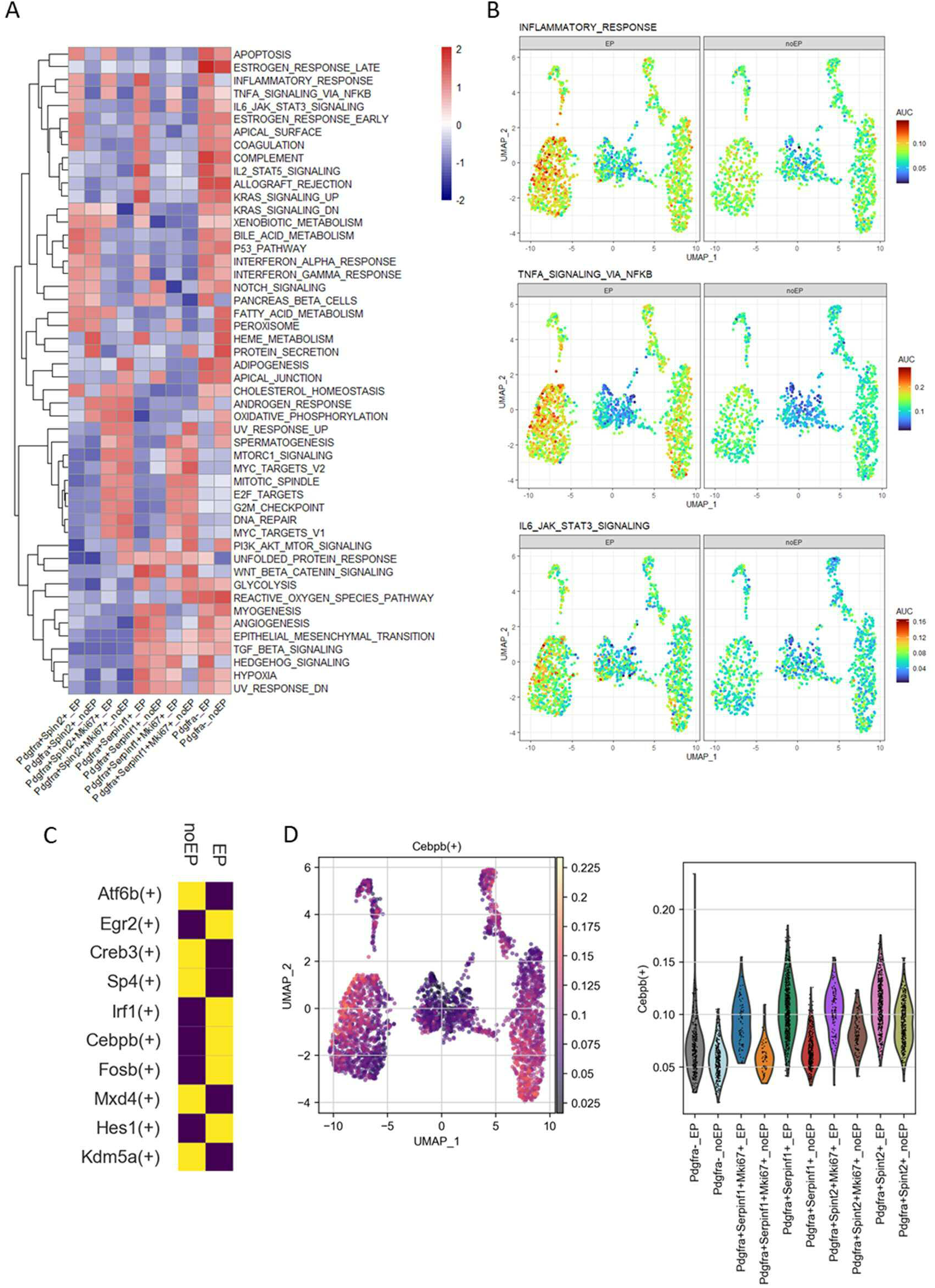
Presence of epithelial cells induce gene expression of TNFA/NFKB and IL-6/STAT3 pathways in fibroblasts. Differences in pathway activities scored per cell type by **(A)** GSVA and individual cell by **(B)** AUCell. **(C)** Heatmap of the binarized regulon activity scores with difference between noEP versus EP group (Purple = no activity, yellow = high activity). **(D)** Activities of Cebpb regulon showed by UMAP and Violin plots.

### Co-culture with epithelium leads to increased expression of IL-6 in fibroblasts

Using differentially expressed gene (DEG) analysis, we found that co-culture with epithelial cells resulted in an increased expression of inflammation related genes in fibroblasts, such as *Nr4a1* (1.4-log2 fold change), *Nfkbia* (1.3-log2 fold change), *Il6* (1.4-log2 fold change) and *Cxcl1* (1.8-log2 fold change) (Fig. 10A). *Il6* was highly expressed in *Pdgfra^+^Serinf1^+^* and *Pdgfra^+^Serinf1^+^Mki67^+^* clusters, while other inflammatory genes were expressed in all clusters (Fig. 10B). The release of IL-6 was significantly increased in co-cultures (Fig. 10C).

**Fig. 10.**
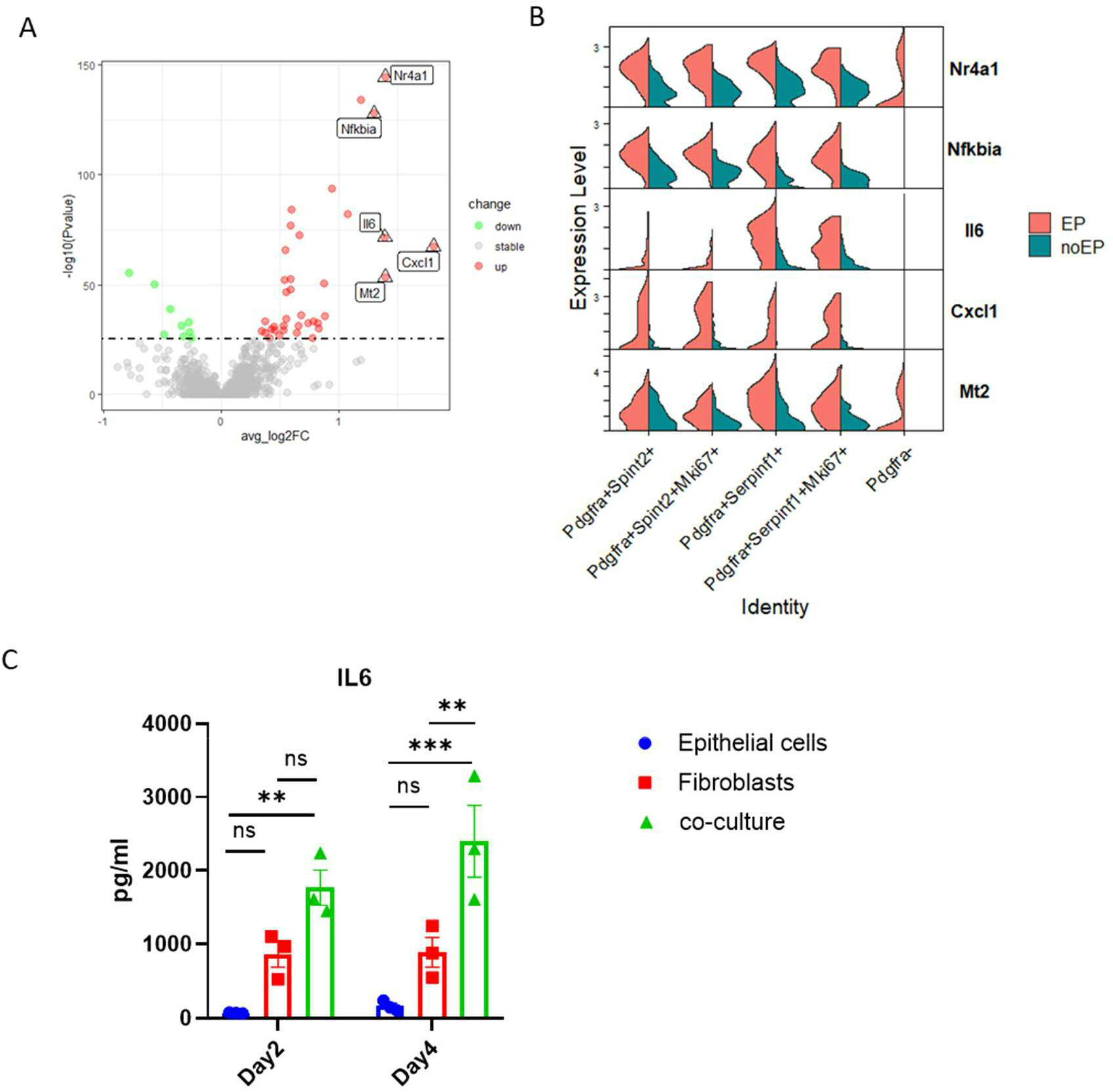
Epithelial cells promote IL-6 expression in fibroblasts. **(A)** Volcano plot showing DEGs of fibroblasts co-cultured with epithelial cells as compared to fibroblast cultured in the absence of epithelial cells. **(B)** Violin plots showing the expression levels of differentially expressed genes in fibroblast sub-clusters. **(C)** Concentrations of IL-6 in supernatants of different ALI culture conditions (n=3 independent experiments). Data are shown as mean ± SEM. Data were compared by two-way ANOVA. **p < 0.01, and ***p < 0.001

Thus, we treated ALI cultures in the absence of fibroblasts with IL-6 resulted in an early increase in transepithelial resistance (Fig. 11A) and expression in *Sftpc, Sftpb* and *Scgb1a1* (Fig. 11B). IF analysis of cytospins showed that IL-6 increased the proportion of club cells expressing HOPX, but not Ki67 positive cells (Fig. 11 C and D).

**Fig. 11.**
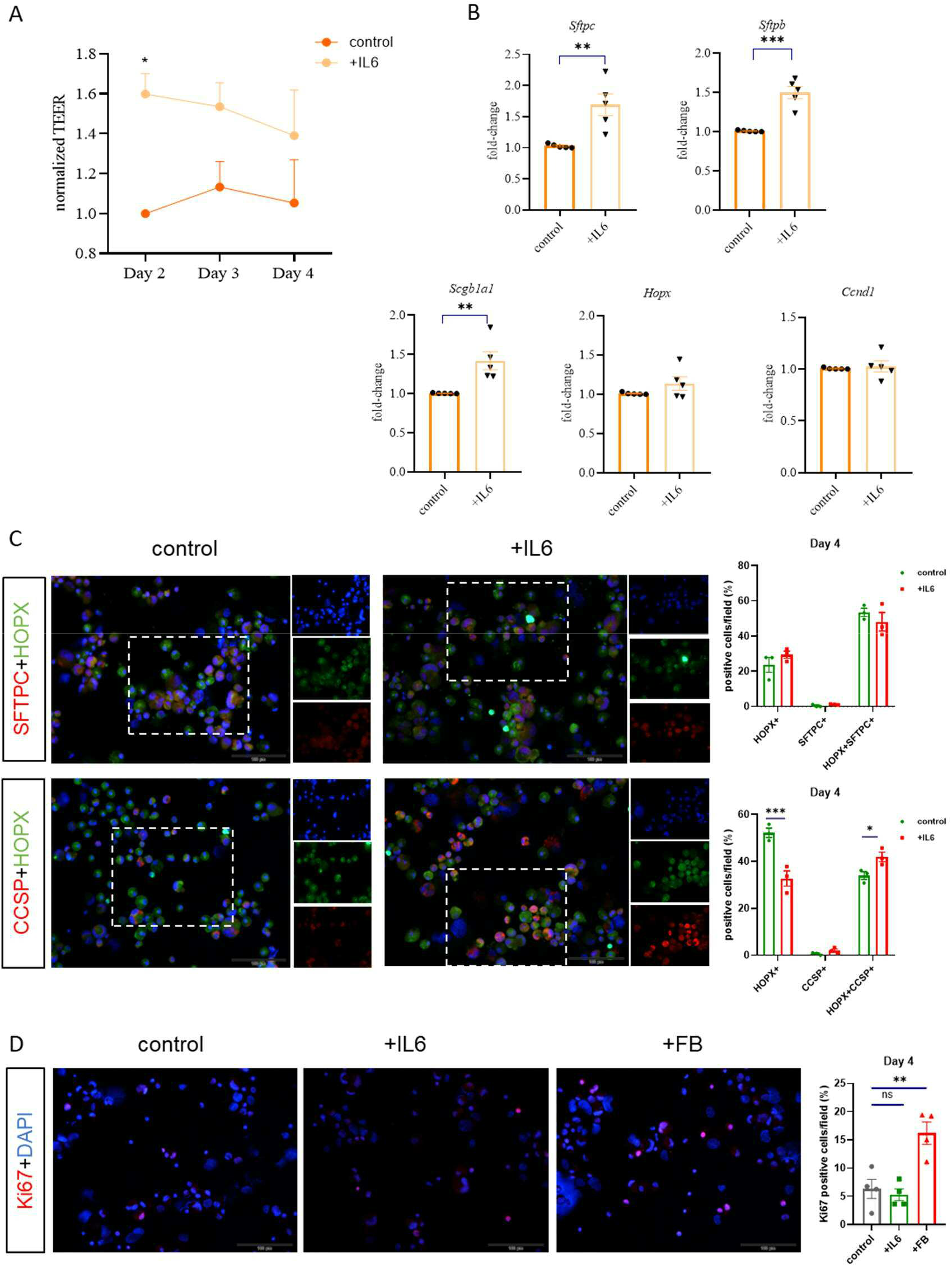
IL-6 promotes barrier formation in ALI cultures without fibroblasts. **(A)** TEER was measured every 24 hours (n=5 independent experiments). **(B)** The expression of selected cell markers as measured by semi-quantitative RT-PCR after 4 days culture with IL-6 (n=5 independent experiments). Cytospins were prepared after four days of culture and SFTPC, HOPX, CCSP **(C)** and Ki67 **(D)** were stained by immunofluorescence. The fraction of cells positive for the different markers was quantified. Data of a representative experiment. Data are shown as mean ± SEM. Data were compared by two-way ANOVA and one-way ANOVA. *p < 0.05, **p < 0.01, and ***p < 0.001.

## Discussion

In this work, we analyzed the interaction of fibroblasts with lung epithelial cells. We identified a regulatory loop, in which AT2 cells interact with the mesenchymal niche in a reciprocal manner to regulate differentiation of both compartments. scRNA-seq analysis showed that in our regeneration model the epithelial barrier is mainly built up by pneumocytes and club cells, with the pneumocytes forming a continuum from AT2 cells via immature AT1 cells to AT1. Co-culture with fibroblasts resulted in an increased epithelial barrier, expression of type 2 markers in pneumocytes, increased activation of regulons mediating AT2 differentiation, and trans-differentiation of club cells towards pneumocytes. Fibroblasts in co-culture increasingly expressed factors that mediate epithelial regeneration.

Co-culture with fibroblasts resulted in increased expression of AT2 markers (e.g. *Sftpb, Lamp3, Cbr2* and *Cxcl15*) [5, 25], in both pneumocyte and club cell clusters. In addition, there was increased activity of the Etv5 regulon in the AT2 and in the club cell cluster, but not in the AT1 cluster. Etv5 has been shown to be essential for the maintenance of AT2 cells. Zhang et al. reported that Etv5 is expressed in AT2 cells in the lung and that deletion of Etv5 in AT2 cells leads to reduced expression of AT2 markers and increased expression of AT1 markers ex vivo [20]. Thus, co-culture with fibroblasts leads to the activation of cellular programs that support AT2 differentiation and drive club cells toward AT2 cells.

On the fibroblast side, co-culture with epithelial cells led to a further increased expression of the growth factors such as FGF7, FGF10 and EREG [23] in the Pdgfra^+^/Serpinf1^+^ clusters. PDGFRα is widely expressed by alveolar and adventitial fibroblasts [11] and PDGFRα^+^ lung stromal cells have been shown two support the growth and differentiation of the alveolospheres cultured in Matrigel [22]. Studies in mice further suggest that the interaction of AT2 cells with *Pdgfra^+^* mesenchymal cells is important in lung repair, but could also be deleterious in lung fibrosis [15, 26, 27]. A cocktail of growth factors released by *Pdgfra^+^* fibroblasts therefore appears to contribute to the improved epithelial barrier and maintenance of AT2 cells in our model. IL-6 has been described to be important for epithelial regeneration [15, 28]. In our model, the co-cultivation with epithelial cells also led to an increased expression of IL-6 in fibroblasts. Moreover, cultivation of epithelial cells in the presence of recombinant IL-6 resulted in an increased epithelial barrier and expression of AT2 markers. These results are consistent with studies showing that IL-6, which is likely released by fibroblasts, can promote AT2 cell growth and self-renewal [15, 29]. IL-6 therefore appears to be a factor supporting the maintenance of AT2 cells in our model. Further studies are needed to clarify to what extent IL-6 regulates AT2 cell maintenance and AT2 differentiation in AT1 cells.

Immunofluorescence staining showed that the vast majority of SFTPC-positive cells were negative for HOPX, a marker associated with AT2 regeneration and AT1 differentiation, immediately after isolation [16, 17]. The isolated pneumocytes are therefore mainly AT2 cells. In contrast, after four days on transwells, most of the SFTPC^+^ cells were also positive for HOPX. In addition, the trajectory analysis of the pneumocyte cluster and the evaluation of cell type-specific markers showed that AT2 cells differentiate via immature AT1 cells to AT1 cells. Thus, the isolated AT2 cells differentiate to a significant extent towards AT1 cells in our model.

Studies have shown that a certain proportion of AT2 cells are progenitor cells that proliferate and contribute to regeneration when lungs are damaged [4, 30]. The question therefore arises to what extent the majority of the isolated AT2 cells in our model build up the barrier, or only certain progenitor cells among the isolated AT2 cells differentiate in the direction of AT1 cells. Since there were no major differences in the ratio of CCSP-to SFTPC-positive cells directly after isolation and after four days on transwells, the vast majority of the isolated epithelial cells seem to contribute to the formation of the epithelial barrier in our model. In any case, our results underline a broad ability of AT2 cells to contribute to the regeneration of the epithelial barrier and to differentiate into AT1 cells.

Our analyzes also showed that club cells were negative for HOPX immediately after isolation. In contrast, differentiation of ALI cultures was accompanied by the expression of HOPX in a proportion of club cells. scRNA-seq data also showed that club cells partially differentiated into AT2 marker-expressing cells, which was further enhanced in the presence of fibroblasts. These results are consistent with studies showing that club cells can differentiate into AT2-type cells and potentially contribute to lung regeneration, at least in mice [14]. However, co-culture with fibroblasts also resulted in increased expression of *Scgb1a1*. This could also be an indication that a large proportion of club cells express certain alveolar markers during regeneration of the alveolar epithelium, but do not differentiate into “true” pneumocytes.

The interaction of fibroblasts with pneumocytes was also examined in organoids models [25]. In the organoid system, however, there is no possibility for the pneumocytes to grow flat, differentiate into real AT1 cells and build an epithelial barrier. In our eyes, the ALI system has far-reaching advantages, such as: Primary cells from knock out mice can be included, a measurable epithelial barrier forms and the effect of stretch stress can potentially be tested, pathogens can be introduced apically and basolaterally in infection experiments, co-culture experiments can be performed in a compartmentalized manner, (nebulized) drugs can be tested in a physiologically relevant way.

Taken together, our results show that mesenchymal cells and epithelial cells interact reciprocally, leading to activation of signaling pathways and expression of factors mediating coordinated alveolar epithelial regeneration. Our results also underscore a broad ability of AT2 cells and club cells to contribute to epithelial barrier regeneration and to differentiate into AT1 cells. Since the epithelial cells grow in a polarized manner in the ALI system, it is very well suited for infection experiments and drug testing.

## Methods

### Isolation of mouse lung distal epithelial cells and fibroblasts

Epithelial cells were obtained from mouse distal lung tissues of C57BL/6 adult mice as described earlier [31, 32]. Tissues were treated with 2 ml dispase (Corning, 354235) for 45 min at 37°C and cut into small pieces and incubated at 37°C for 30-45 min. The cell suspension was filtered through 70 μm and 40 μm cell strainers (Falcon, 352350/352340), and transferred to a culture dish for 90 min at 37 °C to remove fast adhering cells like macrophages. Suspensions were incubated with EPCAM-PE (eBioscience, 12-5791-83, 3 μl/mouse) and anti-PE microbeads (Miltenyi, 130-048-801, 1:20) and cells sorted by MACS. The EPCAM^-^ cells were cultured in DMEM (Gibco, 41965039) for 1~2 passages and stored at −80°C for further using.

### ALI culture for mouse distal epithelial cells

Transwell inserts (6.5 mm; Corning, CLS3470-48EA) were coated with fibronectin (Corning, 354008) and laminin (Sigma-Aldrich, L2020) overnight. 5×10^5^ EPCAM^+^ cells in 200 μl CMM medium based on DMEM/F12 (Gibco, 11320033) supplemented with 1 mM L-glutamine (Life technologies, 25030-081), 10 mM HEPES (Thermo Fisher Scientific, 15630080), 0.25% BSA (Sigma-Aldrich, A9647), 1% NEAA (Sigma-Aldrich, M7145), 100 μg/ml primocin (Invivogen, Ant-pm-1), 0.05% insulin-transferrin-sodium-selenite (Roche, 11074547001) and 10% FBS (Gibco, 10270106) were seeded on transwell membranes. 500 μl CMM with/without 5×10^4^ mesenchymal cells was added to the basolateral compartment. After 48 hours, the apical medium was removed and only medium of the basolateral compartment was refreshed every 2 days. IL-6 at 200 pg/ml (R&D system, 406-ML) was added to the basolateral compartment from day 1, and medium was changed every day.

### TEER. measurement

Millicell^®^-ERS Voltohmmeter (MilliporeSigma, MERS00001) was used to measure TEER. 200 μl prewarmed CMM was added to the Transwell insert. The formula of TEER calculation was as follows: TEER= (the measured resistance – blank resistance) Ω* 0.33cm^2^ (for a 24-well Transwell).

### qPCR

RNA was isolated using the NucleoSpin RNA Kit (Macherey-Nagel, 740955.250) from two biological replicates for each experimental group. The RevertAid RT Reverse Transcription Kit (Thermo Scientific, 1691) was used to generate cDNA. SensiMix™ SYBR^®^ & Fluorescein (2X) reagent (Bioline, QT615-05) was used for all RT-PCR reactions. Samples were placed in a CFX 96^TM^ Real-Time PCR Detection System (Bio-Rad) for 40 cycles running. Threshold cycle values (Ct) for duplicate samples were averaged and normalized to Gapdh (ΔCt), and these values across samples were compared (ΔΔCt) to quantify relative expression. The sequences for the primers were obtained from Origene and primers were obtained from Metabion. Primers are listed in table S1.

### IF staining

Cytospins and ALI Transwell membranes were fixed with 4% paraformaldehyde for 1 hour. After washing with PBS three times, cytospins and membranes were permeabilized by 0.1% Triton X-100 (Roche, SIG10789704001) for 15 min and blocked with 2% BSA in PBS for 1 hour at room temperature (RT). Cells were incubated by anti-pro-SFTPC (Abcam, ab90716, 1:100), anti-HOPX (Santa Cruz Biotechnology, sc-398703, 1:100) and anti-CCSP (Merck, 07-623, 1:100) at 4°C overnight. Secondary goat anti-mouse antibody Alexa 488 (Cell Signaling Technology, 4408S, 1:200) and goat anti-rabbit Cyanine 5 (Invitrogen, A10523, 1:200) were applied for 2 hours at RT in the dark. Cytospins were visualized with an Olympus fluorescent microscope (BX 51, Japan), and membranes were visualized with a Zeiss confocal microscope (LSM900, Germany).

### ELISA

Concentration of IL-6 cytokine in supernatant was quantified using ELISA kit (DuoSet ELISA, DY406) and a BMG microplate reader (Fluostar omega, Germany).

### scRNA-seq

ALI cultures of three Transwells per group were dissociated to obtain single cell solution after 2 days after airlift using TrypLE™ Express Enzyme (Gibco, 12605010). Resulting cell suspensions were washed twice in PBS, counted and assessed for single cell separation and overall cell viability. Cell capture and library preparation were performed using the BD Rhapsody™ Single-Cell Analysis System and the BD Rhapsody™ Whole Transcriptome Analysis (WTA) Amplification Kit (Cat. 633801). For more details see Supplement. The samples were pooled with the BD™ Mouse Immune Single-Cell Multiplexing Kit (Cat. 633793). Cells were captured and final libraries were sequenced on the Novaseq 6000 platform (Illimina, USA) with 50,000 reads per cell. Raw sequencing reads were processed with the BD Rhapsody™ WTA Analysis Pipeline on the Sevenbridges platform.

Unique molecular identifier (UMI) counts were processed with the Seurat package (version 4.1.1) in R software (version 4.1.3). Low-quality cells were filtered out with gene expression <300 genes or the percent of mitochondrial reads over 20% of total reads per cells. The filtered dataset was normalized and scaled by using Seurat NormalizeData (scale factor 10000) and ScaleData function with default parameters. Cell clusters were identified using a shared nearest neighbors (SNN)-based algorithm (resolution was set to 0.4). Nonlinear dimensional reduction was performend to generate UMAP plots as illustrated. Data were integrated using RunHarmony command and cell clusters were identified using a shared nearest neighbors (SNN)-based algorithm. UMAP rendering was performed to visualize the clusters.

Single-cell pseudotime trajectories were generated with the Monocle3 package (Version 1.2.7) in R. FindMarkers function in Seurat was used for DEG analysis. GSVA was used to assess the relative pathway activities in different subtypes of fibroblasts. AUCell package (Version 1.20.2) in R was used to score individual cells for pathway activities. The 50 hallmark gene sets list for signaling pathway activities were derived from the MSigDB databases.

Regulon analysis was done using the python implementation of SCENIC (DOI: 10.1038/nmeth.4463) pySCENIC (version 0.12.0, https://github.com/aertslab/pySCENIC) based on CPM normalized counts. Adjacencies were calculated using pyscenic grn and motifs were pruned with pyscenic ctx against mm10 refseq-r80 10kb_up_and_down_tss.mc9nr.genes_vs_motifs.rankings.feather and mm10 refseq r80 500bp_up_and_100bp_down_tss.mc9nr.genes_vs_motifs.rankings.feather databases. Then pyscenic AUCell was used to obtain regulon activity scores which were visualized using scanpy (version 1.9.1, DOI:10.1186/s13059-017-1382-0).

### Statistics

Graph Prism software (GraphPad Software (version 8.0), San Diego, CA) was used for statistical analysis. Comparisons between groups were analyzed by twoway ANOVA, one-way ANOVA or unpaired t-test as indicated in the figure legends. The results were considered statistically significant for p < 0.05.

## Study approval

Organ harvesting was approved by the Landesamt für Soziales, Gesundheit und Verbraucherschutz of the State of Saarland, Germany in agreement with the national guidelines for animal treatment.

## Author contributions

Y.Y.; F.R.: designing research study, conducting experiments, acquiring data, analyzing data, providing reagents, writing the manuscript; R.B., C.B.: designing research study, analyzing data, providing reagents, writing the manuscript: K.K.: analyzing data; S.M., H.G.: conducting experiments, acquiring data, analyzing data, providing reagents, writing the manuscript; N.SD: designing research study, writing the manuscript.

## Acknowledgments

The authors thank Anja Honecker and Andreas Kamyschnikow (Saarland University, Department of Internal Medicine V) for excellent technical support. This study was supported by grants from Dr. Rolf M. Schwiete Stiftung to RB and Eduard Kastner (Stiftung Forschung für Leben).

