## Supplemental Data for "Mutual regulation of transcriptomes between pneumocytes and fibroblasts mediates alveolar regeneration"

### Supplemental results

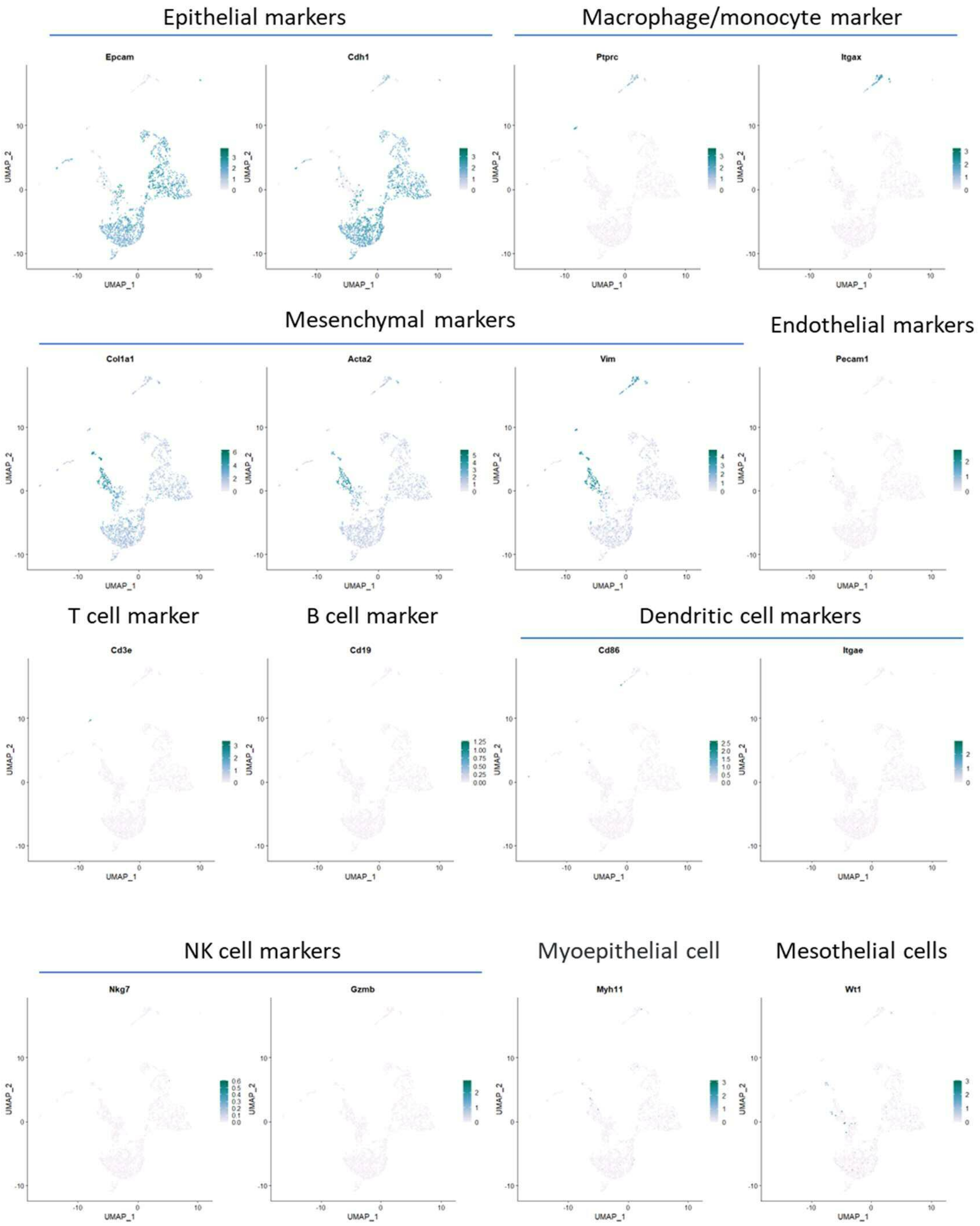


**Supplementary Fig. 1.** scRNA-seq analysis of murine lung cells cultured in ALI. Mouse EpCAM$^{+}$ cells were isolated and cultured on Transwells for four days. UMAP visualization by cell-type specific marker expression.

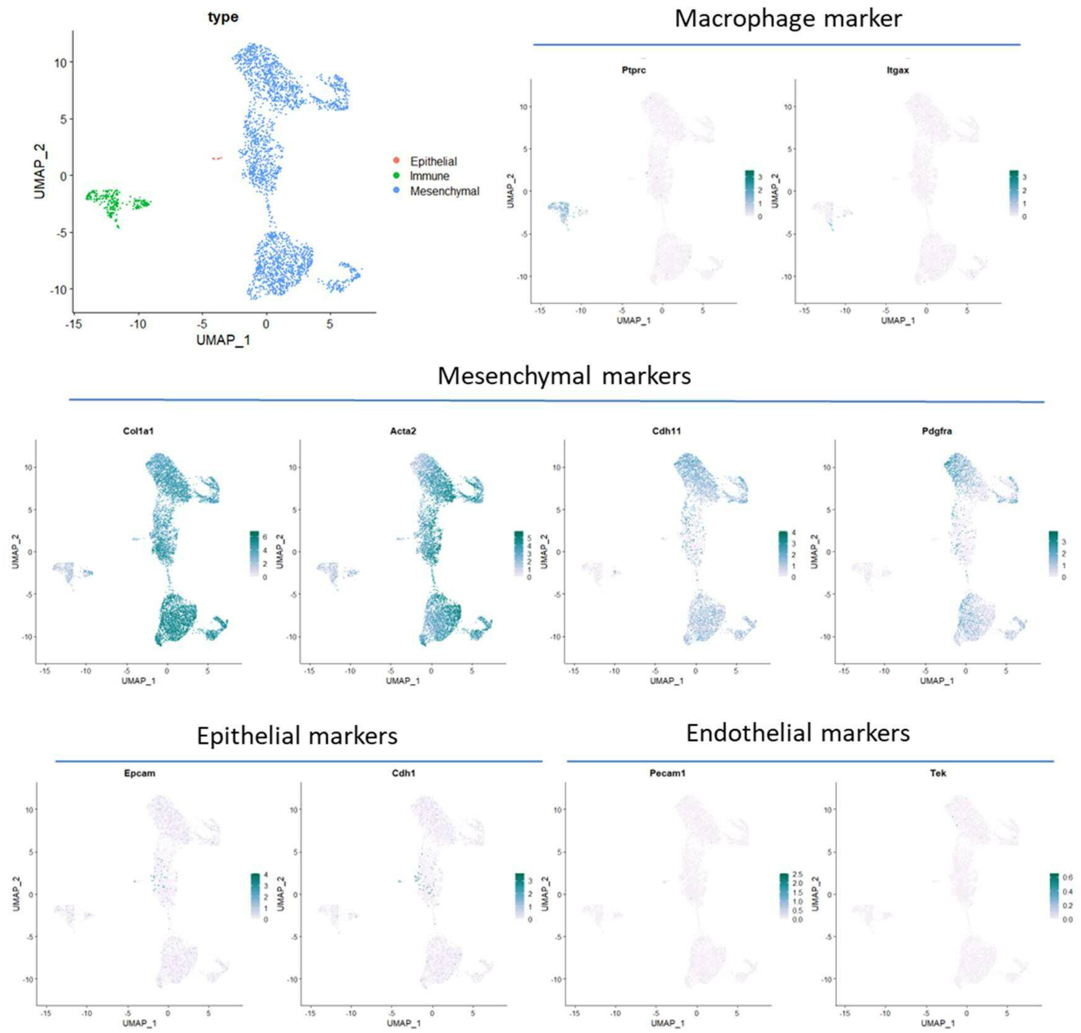


**Supplementary Fig. 2.** scRNA-seq analysis of murine fibroblasts. Mouse EpCAM$^-$ cells were isolated and cultured at the bottom of ALI cultures for four days. **(A)** UMAP clustering of major cell types. **(B)** UMAP visualization of cell type-specific markers expression.

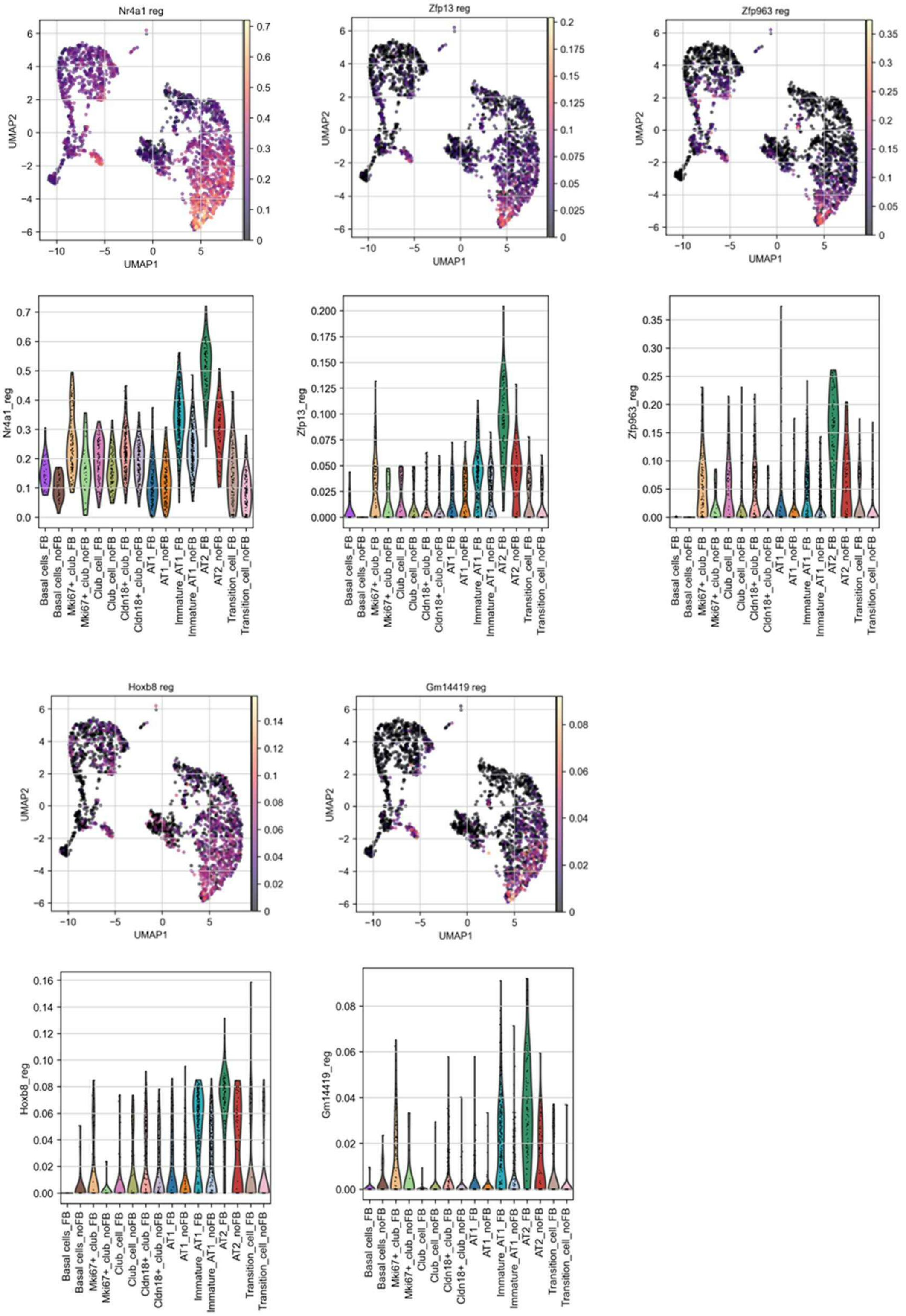

Nr4a1 reg
Zfp13 reg
Zfp963 reg
Hoxb8 reg
Gm14419 reg
UMAP1
UMAP2
Nr4a1_reg
Zfp13_reg
Zfp963_reg
Hoxb8_reg
Gm14419_reg
Basal cells_FB
Basal cells_noFB
Mki67+_club_FB
Mki67+_club_noFB
Club_cell_FB
Club_cell_noFB
Cldn18+_club_FB
Cldn18+_club_noFB
AT1_FB
AT1_noFB
Immature_AT1_FB
Immature_AT1_noFB
AT2_FB
AT2_noFB
Transition_cell_FB
Transition_cell_noFB

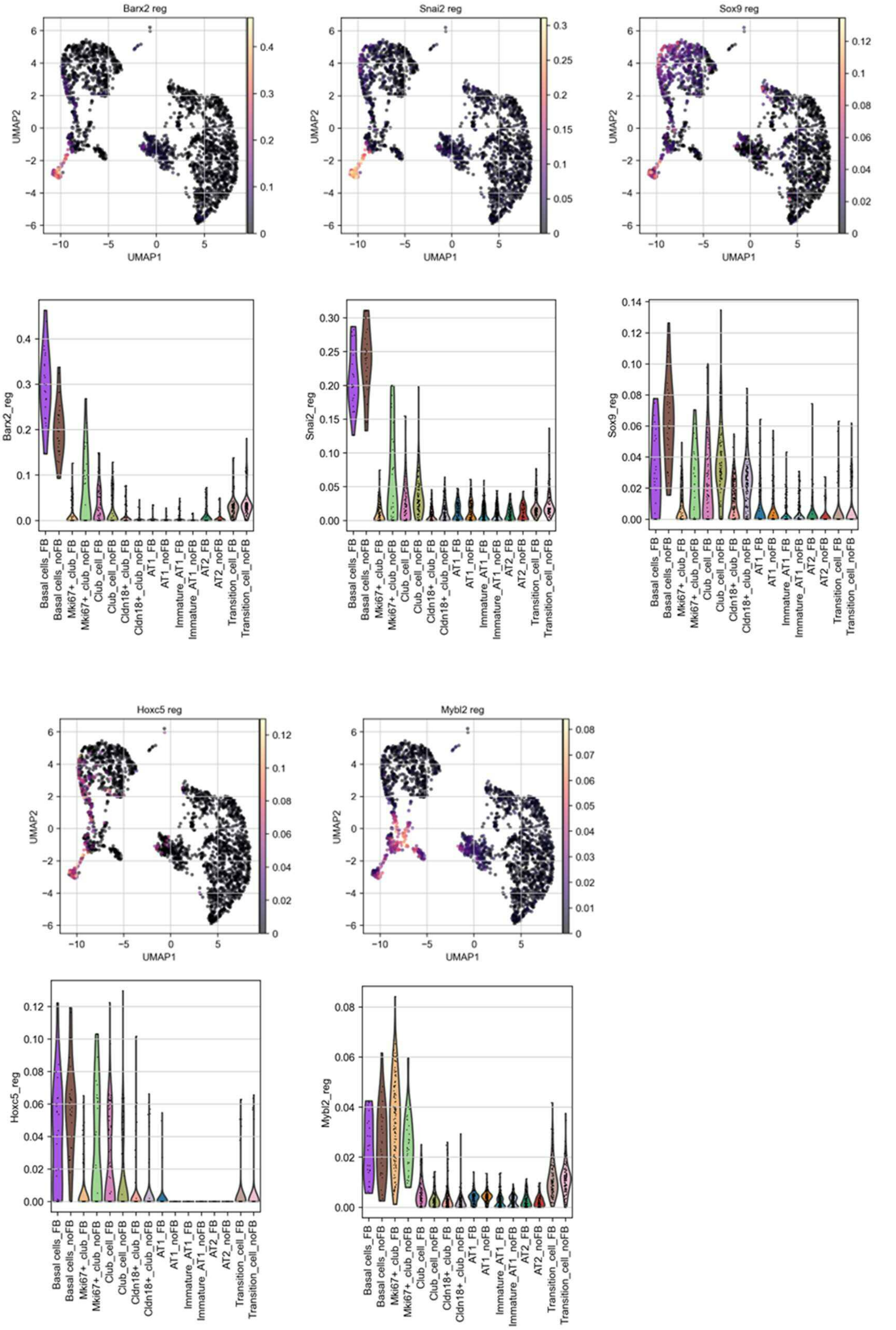


**Supplementary Fig. 3.** Regulon analysis in epithelial cells. UMAP representation and corresponding violin plots of regulon activity.

**Table 1:** RT-PCR primer sequences

| Primer | Sequence |
|---|---|
| Mouse *Scgb1a1*[1] | Forward: 5’-GGTTATGTGGCATCCCTGAAGC-3’<br>Reverse: 5’-GCTTACACAGAGGACTTGTTAGG-3’ |
| Mouse *Sftpc* | Forward: 5’-GTCCTCGTTGTCGTGGTGATTG-3’<br>Reverse: 5’-AAGGTAGCGATGGTGTCTGCTC-3’ |
| Mouse *Sftpb* | Forward: 5’- TGTCCTCCGATGTTCCACTGAG-3’<br>Reverse: 5’- AGCCTGTTCACTGGTGTTCCAG-3’ |
| Mouse *Aqp5* | Forward: 5’- TCCATGAACCCAGCCCGATCTT-3’<br>Reverse: 5’- GAAGTAGAGGATTGCAGCCAGG-3’ |
| Mouse *Hopx* | Forward: 5’- TCTCCATCCTTAGTCAGACGC-3’<br>Reverse: 5’- GGGTGCTTGTTGACCTTGTT-3’ |
| Mouse *Ccnd1* | Forward: 5’- GCAGAAGGAGATTGTGCCATCC-3’<br>Reverse: 5’- AGGAAGCGGTCCAGGTAGTTCA-3’ |
| Mouse *Gapdh* | Forward: 5’- CATCACTGCCACCCAGAAGACTG-3’<br>Reverse: 5’- ATGCCAGTGAGCTTCCCGTTCAG-3’ |

### Supplemental methods

#### Single cell sequencing analysis

Single cell analysis was performed using the BD Rhapsody™ Single-Cell Analysis System according to manufacturer’s protocols. Thus, if not stated otherwise reagents were obtained from Becton Dickinson (Heidelberg, Germany). For multiplex purposes, cells of each experimental condition were initially incubated with an individual barcode labelled antibody (Mouse Single Cell Sample Multiplexing Kit, Cat. No. 626545) for 20 minutes at room temperature. Cells were washed twice with BD Pharmingen Stain Buffer (Cat. No. 554656) and viability-stained with 2 mM Calcein AM (Cat. No. C1430; Thermo Fisher Scientific, Dreieich, Germany) and 0.3 mM Draq7 (Cat. No. 564904) for 5 minutes at 37°C. Cells were counted using a disposable hemocytometer (Cat. No.

DHCN01-5; INCYTO, Cheonan, South Korea) and cell viability was determined. The BD Rhapsody Cartridge (Cat. No. 400000847) was primed with 100% ethanol followed by 2 washes with Cartridge Wash Buffer 1 (Cat. No. 650000060) and one wash with Cartridge Wash Buffer 2 (Cat. No. 650000061). About 30.000 labelled cells were loaded and incubated for 15 minutes at room temperature. Excess fluid was removed, the cartridge was loaded with Cell Capture Beads (Cat. No. 650000089) and incubated for 3 minutes at room temperature. Excess beads were washed off using Sample Buffer. Lysis Buffer was applied and beads were extracted from the cartridge using the BD Rhapsody Express instrument and washed twice with cold Bead Wash Buffer (Cat. No. 650000065). The cDNA reaction mix was prepared as indicated in the manufacturer's protocol and mixed with the beads. The mixture was incubated in a thermomixer (37 °C, 1200 rpm, 20 minutes). The supernatant was removed and replaced by the Exonuclease I mix prepared according to the manufacturer's protocol. The bead suspension was incubated in the thermomixer (37°C, 1200 rpm, 30 minutes, followed by 80 °C without shaking for 20 minutes). The suspension was then briefly placed on ice and the supernatant was removed. Finally, beads were resuspended in Bead Resuspension Buffer (Cat. No. 650000066).

Single cell mRNA and multiplex sample Tag libraries were prepared using the BD Rhapsody WTA Amplification Kit (Cat. No. 633801) according to manufacturer's recommendation (BD Rhapsody system mRNA WTA and Sample Tag library protocol, 23-21712-00). Briefly, the Bead Resuspension Buffer was removed, beads were resuspended in Elution Buffer and incubated on a heat block at 95 °C without shaking for 5 minutes. The tube was briefly centrifuged and the supernatant was retained as Sample Tag product. The Random Primer Mix was prepared as indicated in the manufacturer's protocol and were mixed with the beads and incubated using the following conditions: heat block without shaking at 95 °C for 5 minutes, and at 1200 rpm in the thermomixer at 37°C for 5 minutes followed by 25°C for 15 minutes. The Extension Enzyme Mix was prepared as indicated in the manufacturer's protocol and was added to the beads and incubated at 1200 rpm in the thermomixer at 25°C for 10 minutes, 37°C for 15 minutes, 45°C for 10 minutes, and 55°C for 10 minutes. Primer and Enzyme Mix was replaced by Elution Buffer and denaturated on a heat block without shaking at 95 °C for 5 minutes. Afterwards the beads were resuspended by use of the thermomixer at 1200 rpm for 10 seconds. The supernatant was retained as Random Primer Extension Product (RPE Product) and purified using AMPure XP magnetic beads (Cat. No. A63880; Beckman Coulter, Krefeld, Germany). The purified product was further amplified by a second PCR of 12 cycles and again purified using AMPure XP beads, resulting in the RPE PCR product. The concentration of the RPE PCR product was determined using the Agilent 2100 Bioanalyzer with the High Sensitivity DNA Kit (Cat. No. 5067-4626; Agilent, Waldbronn, Germany). The RPE PCR product was diluted with Elution Buffer to yield concentration of 2 nM of the 150 - 600 bp peak and was then amplified by the final WTA index PCR for 8 cycles with subsequent dual-sided cleanup using AMPure XP beads afterwards.

The Sample Tag PCR1 reaction mix was prepared as indicated in the manufacturer's protocol and Sample Tag PCR1 reaction mix was mixed with Sample Tag product. The Sample Tag product was amplified in the thermal cycler for 11 cycles of the PCR program indicated in the protocol and subsequent purification using AMPure XP beads. Sample Tag PCR1 product was amplified with a second PCR of 10 cycles and subsequent purification using AMPure XP beads. The resulting Sample Tag PCR2 product was amplified by the final Sample Tag index PCR for 7 cycles with subsequent purification. Concentrations of WTA and Sample Tag index PCR products were determined using the Qubit Fluorometer and the Qubit dsDNA HS Kit (Cat. No. Q32851; Thermo Fisher Scientific) and quality control was performed on the Agilent 2100 Bioanalyzer with the High Sensitivity DNA Kit.

VEGF receptor 2 (KDR) protects airways from mucus metaplasia through a Sox9 dependent pathway
